# Prevalence of scabies and its associated factors among school age children in Arba Minch zuria district, Southern Ethiopia, 2018

**DOI:** 10.1101/2020.03.16.993576

**Authors:** Abayneh Tunje, Chuchu Churko, Desta Haftu, Amsalu Alagaw, Eyayou Girma

## Abstract

**Background:** Scabies, a common human skin disease with a prevalence range of 0.2% to 71.4% in the world. It can have considerable impact on general health leading to illness and death not only through direct effect of its infestation and as a result of secondary bacterial infection. The aim of this study was to assess the prevalence of scabies and its associated factors among school age children in Arba Minch zuria district, Gamo zone, Southern Ethiopia.

**Methods:** A community based cross sectional study was carried out in 845 school age children from February 20 to March 30, 2018. Multi-stage sampling technique was used to select study populations. Logistic regression an analysis was used to identify factors associated with scabies. Findings were presented using 95% CI of Crude Odds Ratios (COR) and Adjusted Odds Ratios (AOR. To declare statistical significance, p-value less than 0.05 was used.

**Result:** A total of 825 children participated in the study with response rate of 97.6%. The overall prevalence of scabies was 16.4% [95% CI: 13.9%, 18.9%]. overcrowding index, family history of itching in the past two weeks, wealth index, knowledge of scabies, climatic zone, frequency of washing body, frequency of washing clothes, finger nails cutting practice, history of skin contact with scabies patient, washing hair more than once weekly, and sharing of clothes were significantly associated scabies disease.

**Conclusion:** In conclusion, the prevalence of scabies in Arba Minch lies at 16.4% in the global scabies range 0.2% to 71.4%. The prevalence was highest in highlands followed by midland and then lowland. This represents a significant scabies burden which we recommend warrants health service intervention.

**Authors summary:** Scabies, a common human skin disease with a prevalence range of 0.2% to 71.4% in the world. It can have considerable impact on general health leading to illness and death not only through direct effect of its infestation and as a result of secondary bacterial infection. Conducting a research on this neglected tropical disease would contribute in designing a policies and strategies on prevention and control measures in the study area. Therefore, the aim of this study was to assess the prevalence of scabies and its associated factors among school age children in Arba Minch zuria district, Gamo zone, Southern Ethiopia.

## Introduction

The skin is the largest body organ and is a sensitive indicator of child’s general health. Skin disorders are the most common health problems among children. Skin disorder may cause emotional and psychological stress for the child and family [1]. Scabies is contagious skin infestation caused by the parasite Sarcoptes scabiei. It is the most common ectoparasitic dermatosis seen in clinical practices [2].

The International Alliance for the Control of Scabies, a newly formed organization, proposes to achieve scabies control in vulnerable communities in 2013 [3]. Recently in the tenth strategic and technical advisory group for neglected tropical diseases World Health Organization (WHO) added scabies to the list of “Neglected Tropical Diseases”, thereby recognizing its impact on human health [4].

Ethiopia has been affected by natural disasters such as the El Niño weather phenomenon, leading to severe drought and scabies outbreaks [5]. The country faced scabies outbreak in many parts of the regions due to this weather event. [5]. Ethiopia has developed a guideline to assist with control and to attempt prevent scabies outbreaks in response to these outbreaks. The recommended interventions include: Health, Water, Sanitation and Hygiene (WASH) and communication for development. Considering scabies as water washed disease, the key intervention is the provision of access to sufficient safe water for personal hygiene: Washing of clothing, washing of body using soaps especially the affected areas, appropriate hand washing at critical times, clothing or bedding that were used by an infested individual during and before effective treatment should be dried for 3 days in the sun to allow time for mites and eggs to die. The communication strategies for prevention of scabies need integrated and multi-level interventions comprising advocacy, social mobilization, social and behavioral change communication at different level and capacities should be provided in Ethiopia [6].

Scabies is a common problem in school age children and there is limited study done on scabies in the study area. Therefore, the aim of this study was to assess the prevalence of scabies and its associated factors among school age children in Arba Minch zuria district, Gamo zone Southern Ethiopia.

## Methods

### Study area and period

Community based cross-sectional study was conducted from March 1-30, 2018 in Arba Minch Zuria district, Gamo Gofa Zone. The distrct is located 505 kilometers south of Addis Ababa, capital city of Ethiopia. The district has 29 kebeles (the lowest administrative structure in Ethiopia, similar to village). According to the central statistical agency of Ethiopia in 2007 the total population of the district was 165,680 of which 82,751 were males and 82,929 were females. The district has 7 health centers and 40 health posts.

### Population

The source populations were all children aged 5-14 years in the selected rural kebeles who fulfill selection criteria during the study period. All children aged 5 to 14 years in the selected kebeles were included in the study, and children who severely ill were excluded from the study.

#### Sample size and sampling

The sample size was determined using single population proportion formula with the assumption of 95 % confidence interval (CI), by taking prevalence of scabies 50%, design effect [DE] = 2, none response rate of 10%. The final, the calculated sample size was 845. The study participants were recruited using multi-stage sampling technique.

#### Data collection

Socio-demographic characteristics and associated risk factors were collected using structured questionnaire by trained health professionals. Structured questionnaire was used to interview Parents and children in their home. Physical examination was undertaken on respondents who had itchy vesicular skin rash by the interviewers. The diagnosis was ascertained, based on the Mali clinical algorithm[7]. Skin scraping was not feasible in this setting.

### Operational definitions

#### Scabies

in this study scabies is defined as the presence of persistent pruritic rash with itching increasing at night which are notified at least at two specific body sites(on the wrist, sides and web spaces of the fingers, the axillae, periareolar, per umbilical, genitalia area, abdomen, and buttock areas) with or without history of pruritus in the close entourage [7].

#### Good knowledge

Those mother/care giver who answered above the mean of the knowledge questions.

#### Poor knowledge

Those mother/care giver who answered below the mean of the knowledge questions.

#### School age children

children who were in the age group 5-14 years old [8].

#### Infrequent bathing

showering frequency less than once per week in the past one month [9].

#### Infrequent washing clothes

washing clothes less than once per week in the past one month [9].

#### Infrequent changing clothes

changing clothes less than once per week in the past one month [9].

#### Overcrowding index

was calculated by dividing the number of usual residents in a house by the number of bedrooms in the house. If it is more than 1.5 overcrowded and if it is less than or equal to 1.5 not overcrowded [10].

### Data quality control

To ensure quality of data, questionnaire was prepared in English language, translated to Amharic and re translated back to English by other person who can speak both languages. To make sure that the questionnaire is appropriate and understandable; it was pre-tested on 5% of sample size.

Training was given for supervisors and data collectors for one day. Regular supervision was carried out during data collection period’. ‘Collected data were checked for completeness and consistency daily.

### Data analysis

Epi info version 7 was used for entering, coding, cleaning the collected data that were analyzed using SPSS version 20. In the univariate analysis a descriptive statistics was conducted to explore frequency distribution, central tendency, variability (dispersion) and overall distribution of independent variables. Bivariate analysis was done to determine the associations between each independent variable and outcome variable. All associated factors with p-value less than during bivariate analysis and biologically plausible factors were entered in to multivariable logistic regression model. Odds ratio with 95% confidence intervals was used to see the strength of association between different variables. P-value and 95% confidence interval (CI) for odds ratio (OR) were used in deciding the significance of the associations. Wealth index was calculated by using principal component analysis method (PCA) and constructed as lowest, second, middle, fourth and highest.

### Ethical considerations

Ethical approval was obtained from Arba Minch University, Institute Review Committee (IRC of AMU, Ref.No. 10994/111). Permission letters were obtained from the woreda administration office and the selected kebele leaders. The childrens’ families were informed about the objective of the study and written consent and assent was obtained. The respondent’s confidentiality was maintained. Those children and families who suffered from scabies and who developed secondary complication were referred to health facility for anti-scabies medication.

## Results

### Socio-demographic characteristics of the respondents

A total of 845 (response rate of 97.6%) school age children were participated in the study. Of the total children 53.2% (439/825) were men, and 54.9% of the children. Majority 81.6% (673/825) of the study subjects attended school whereas 18.4% (152/825) did not. Majority 91.3% of the respondents were Gamo ethnic group, 5.6% were Wolayta, 1.8% Amhara and the rest 1.3% others. More than half (60.1%) were follower of Protestant Christianity followed by Orthodox Christianity (38.8%) and 1.1% others. 50.4% (416/825) of the respondents were from midland, 39.4% (325/825) were lowland and 10.2% (84/825) highland. Regarding family education, 50.0% of the mothers had no formal education, 44.2% had primary education, and 5.8% had secondary and higher educational level. 43.9% of children’s father had no formal education, 41.1% had attend primary education and 15% had secondary and higher educational status. Majority 92.6% (764/825) of children’s mother interviewed in this study were housewife, 5.8% were merchants and 1.6% employed Gov’t/NGO. On the other hand, 91.6% (756/825) of the children’s fathers farmer, 6.3% (52/825) and 2.1% (17/825) were Merchant and employed respectively (table 1).

**Table 1:**
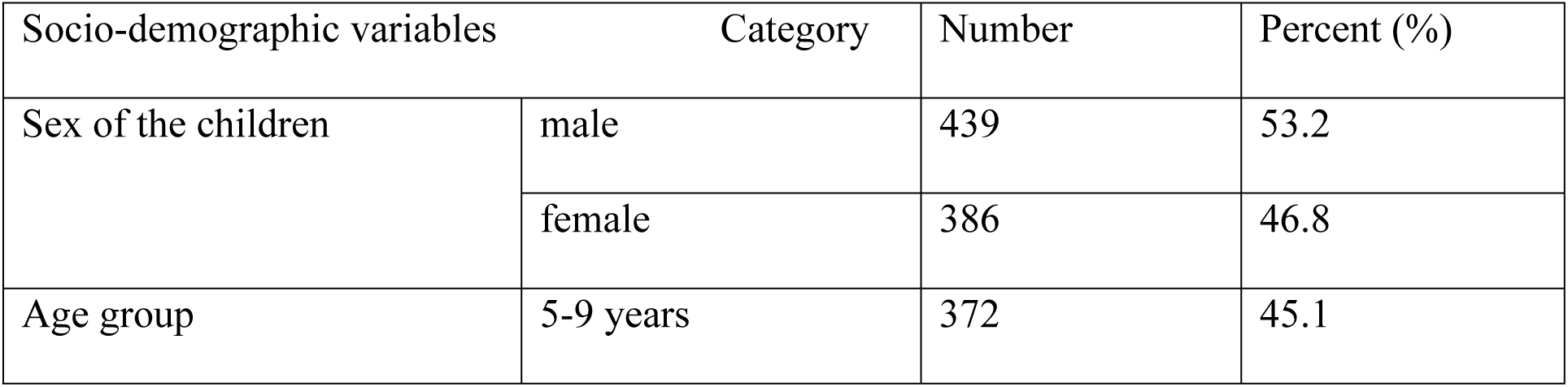
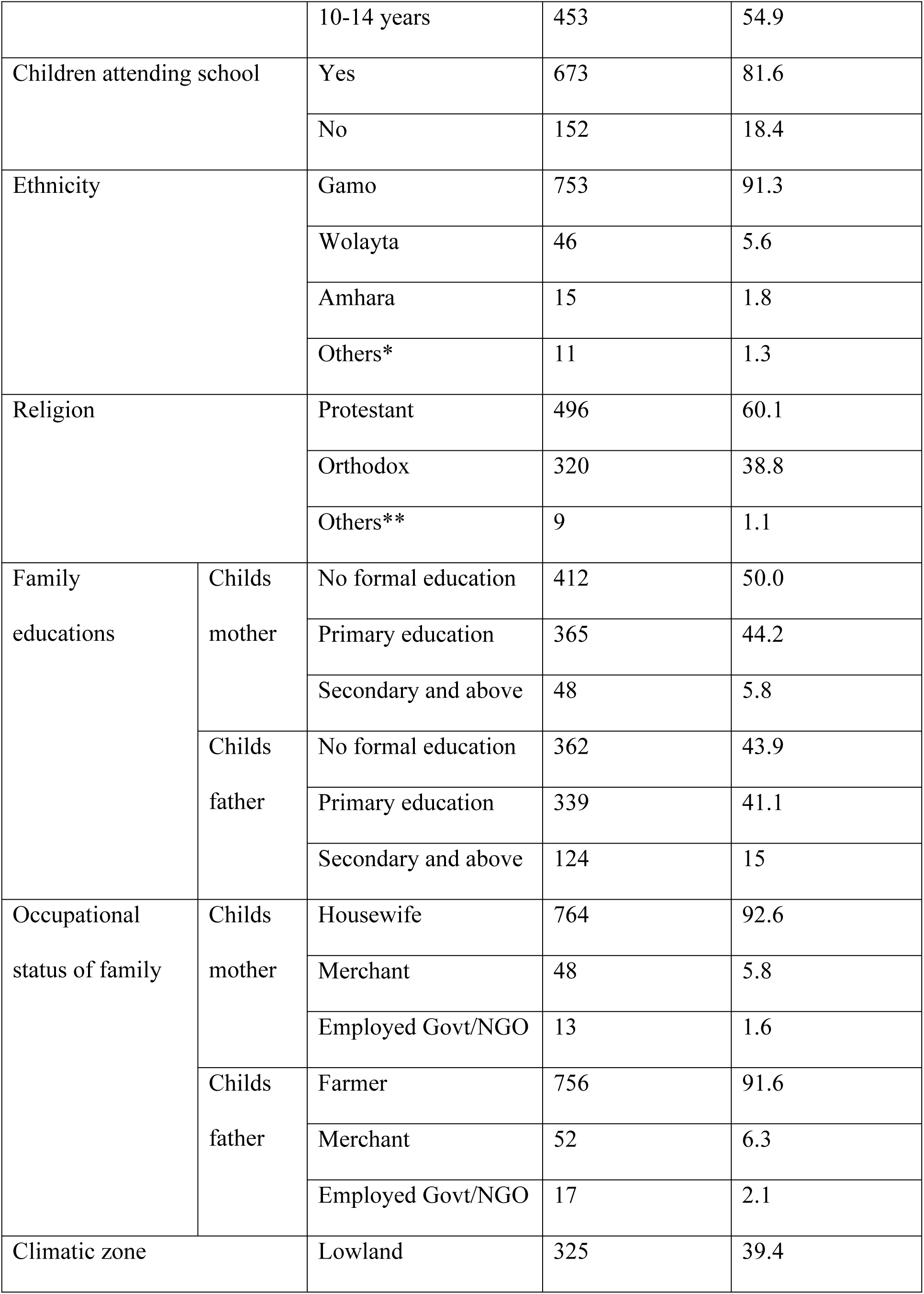
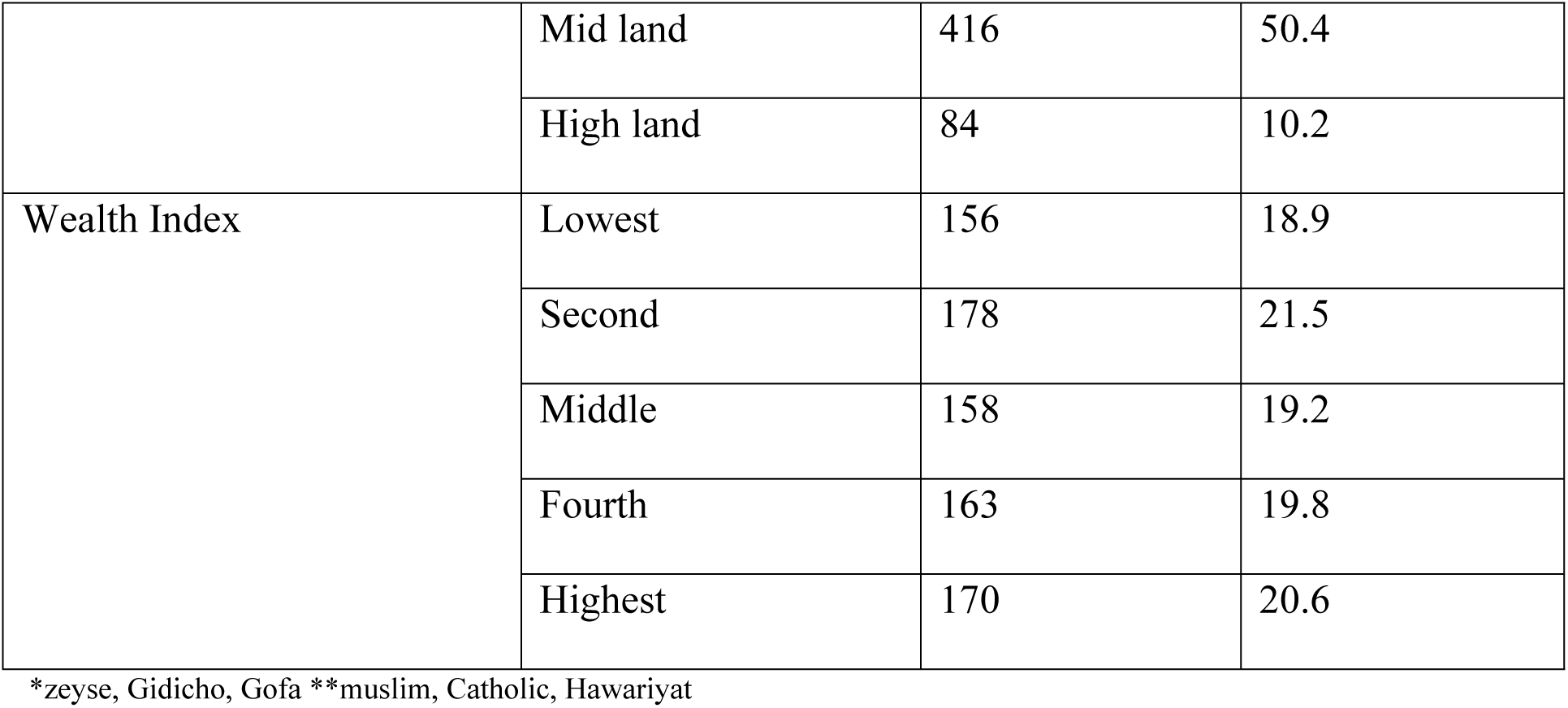
Socio-demographic characteristics of the study subjects in Arba Minch zuria district, SNNPR, 2018.

### Prevalence of scabies among school age children

The overall prevalence of scabies among school age children 16.4% (135/825) [95% CI: 13.9%, 18.9%] scabies cases were identified. Out of which 45 (13.8%) [95%CI: 10.05%, 17.55%] were found in lowland, 65 (15.6%) [12.11%, 19.09%] in midland and 25 (29.8%) [95%CI: 20.019%, 39.58%] found in highland area. The prevalence of scabies among females was higher than male children, 17.1% and 15.7% respectively.

### Home environment related factors

More than three quarters of the sample, 638/825 (77.3%) had family size of greater than or equal to five and only 187/825 (22.7%) had less than five family members with the mean family size of 6 and standard deviation 1.95. 80.2% (662/825) of the children share bed with others whereas 19.8% (163/825) did not share their bed with other family member. From those families who had animals in their home 366/825 (55.5%) of the children look after with animals and 294/825 (44.5%) did not deal with animals in their home. 474/825 (57.5%), 239/825 (28.9%) and 112/825 (13.6%) of the respondents use electricity, kerosene light and solar light in their home respectively. Majority 561 (68%) of the respondents house was covered by iron sheet and 264 (32%) did not. Almost all 815 (98.8%) of the family were living in a house built by soft bricks and 10 (1.2%) living in a hard bricks.

Majority 586/825 (71%) of the respondents use river/pond as a source of water for personal hygiene, 136/825 (16.5%) use pipe/tap water and the rest 103/825 (12.5%) use well/spring water in the study area. Regarding knowledge 636/825 (77.1%) of the family had good knowledge about scabies and 189/825 (22.9%) had poor knowledge. About 747 (90.5%) had water source near their home <30 minutes and 78 (9.5%) had water source far away from home (table 2).

**Table 2:** Home environment factors of the respondents in the study area, 2018.

| Variables | Category | Frequency | Percentages |
| --- | --- | --- | --- |
| Family size | <5 | 187 | 22.7 |
|  | >=5 | 638 | 77.3 |
| Bed sharing | Yes | 662 | 80.2 |
|  | No | 163 | 19.8 |
| Dealing with animals | Yes | 366 | 55.5 |
|  | No | 294 | 44.5 |
| Type of light | Kerosene | 239 | 28.9 |
|  | Solar | 112 | 13.6 |
|  | Electricity | 474 | 57.5 |
| House type iron sheet covered | Yes | 561 | 68 |
|  | No | 264 | 32 |
| Type of house built by | Soft bricks | 815 | 98.8 |
|  | Hard bricks | 10 | 1.2 |
| Source of water for personal hygiene | Pipe/tap water | 136 | 16.5 |
|  | Well/spring | 103 | 12.5 |
|  | River/pond | 586 | 71 |
| Knowledge status | Good | 636 | 77.1 |
|  | Poor | 189 | 22.9 |
| Distance of water source from home | Near home/$\leq 30$ minute | 747 | 90.5 |
| | Far away from home $> 30$ minute | 78 | 9.5 |

Regarding personal hygiene, 85.8% (708/825) of the respondents wash their body frequently and 14.2% (117/825) had responded that they wash their body infrequently. Majority 80.2% (662/825) of the participants wash their clothes frequently, 19.8% (163/825) wash their clothes infrequently. More than three quarters 83.5% (689/825) of the respondents changed into clean clothes frequently whereas the rest 16.5% (136/825) changed clothes infrequently. 56.7% (468/825), 280/825 (34%) and 77/825 (9.3%) of the children wash their hair 1-7 days, 7-14 days and more than 14 days respectively. 627/825 (76%) of the children share their clothes with any other person whereas 198/825 (24%) did not share their clothes. About 563 (68.2%) of the children cut their finger nails short/trimmed and 262/825 (31.8%) did not (table 3).

**Table 3:** personal hygiene and sanitation characteristics of the respondents, 2018.

| Variables | Category | Frequency | Percentages |
| --- | --- | --- | --- |
| Body washing | Frequently | 708 | 85.8 |
|  | Infrequently | 117 | 14.2 |
| Washing clothes | Frequently | 662 | 80.2 |
|  | Infrequently | 163 | 19.8 |
| Changing of clothes | Frequently | 689 | 83.5 |
|  | Infrequently | 136 | 16.5 |
| Frequency of hair washing | 1-7 days | 468 | 56.7 |
|  | 7-14 days | 280 | 34 |
|  | >14 days | 77 | 9.3 |
| Share clothes with other | Yes | 198 | 24 |
|  | No | 627 | 76 |
| Finger nails cut short/trimmed | Yes | 563 | 68.2 |
|  | No | 262 | 31.8 |

### Knowledge about scabies among children’s family

From the total respondents, 805/825 [97.6%] knew the signs and symptoms of scabies, and 78.7% knew parts of the body that are affected by scabies as finger webs, armpits, genitalia, abdomen, breast, waist and knees, 143/825 [17.3%] knew that scabies affects parts of body that are mostly covered and 33 [4%] said it affects mostly at genitalia area (table 4).

**Table 4:** shows sites of scabies rash among children diagnosed scabies infection in the study area, 2018.

| Sites of the rash | Frequency | Percentage |
| --- | --- | --- |
| Finger webs and ulnar area | 135 | 100% |
| Auxiliary area | 59 | 43.7% |
| Genitalia | 45 | 33.3% |
| Abdomen | 104 | 77% |
| Shoulder blades | 7 | 5.2% |
| Elbow | 8 | 5.9% |
| Buttocks | 72 | 53.3% |
| Lower legs | 16 | 11.9% |

Regarding respondents knowledge about transmission, 545/825 [66.1%] knew scabies transmitted through skin to skin contact and infected fomites like clothing, bed linen, 213 [25.8%] through skin to skin contact only, and 67 [8.1%] through fomites (table 5).

**Table 5:**
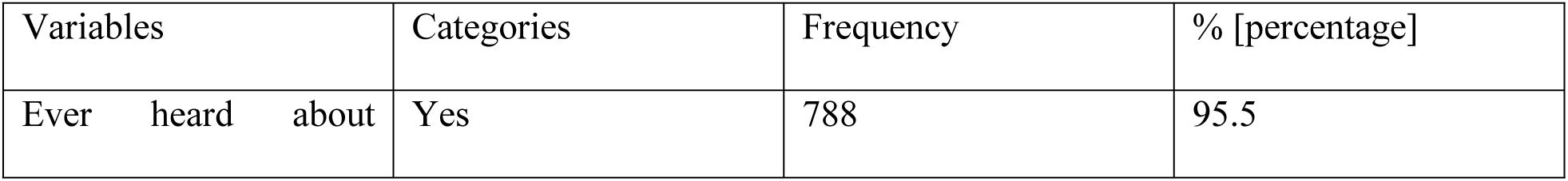
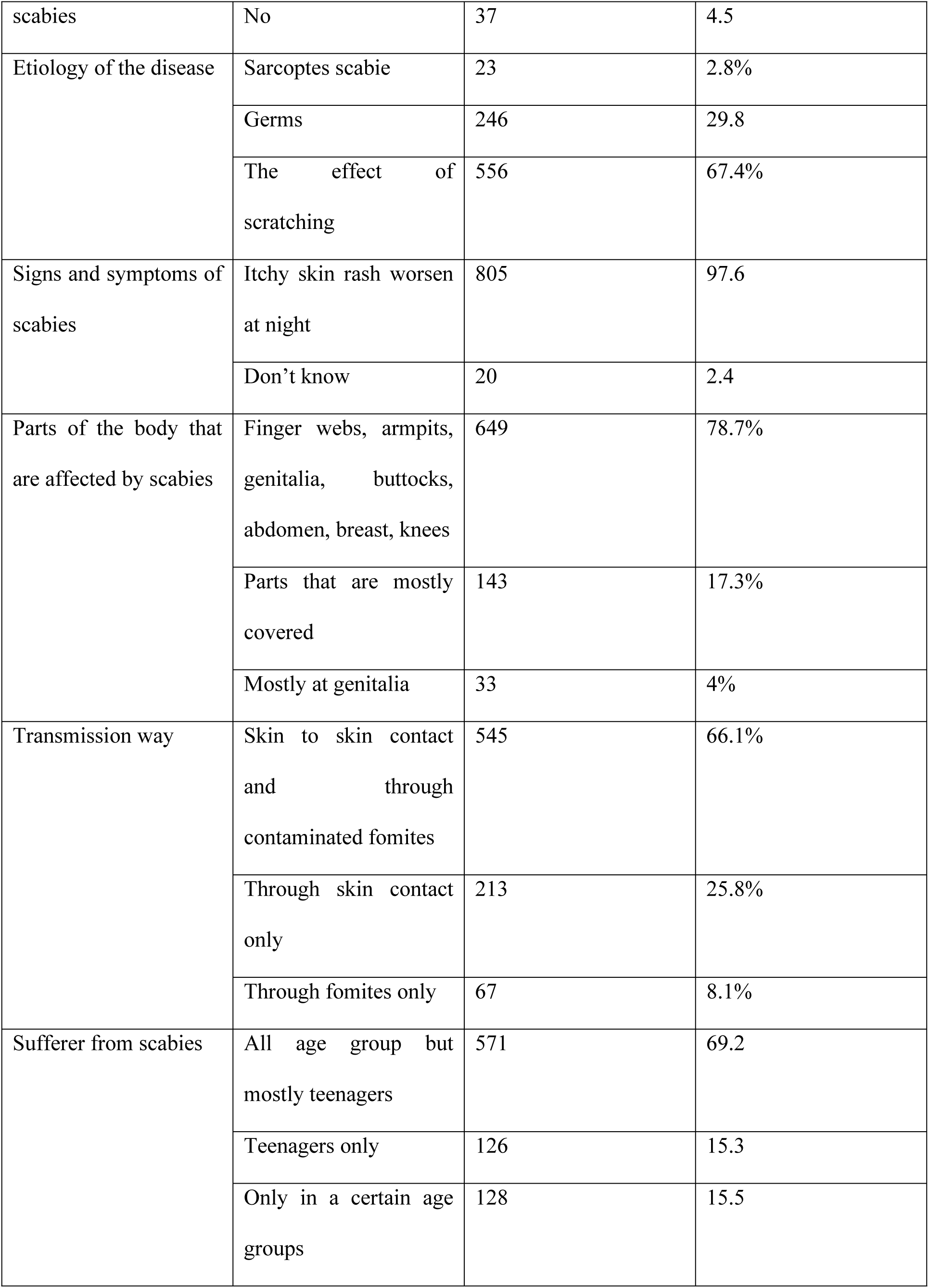
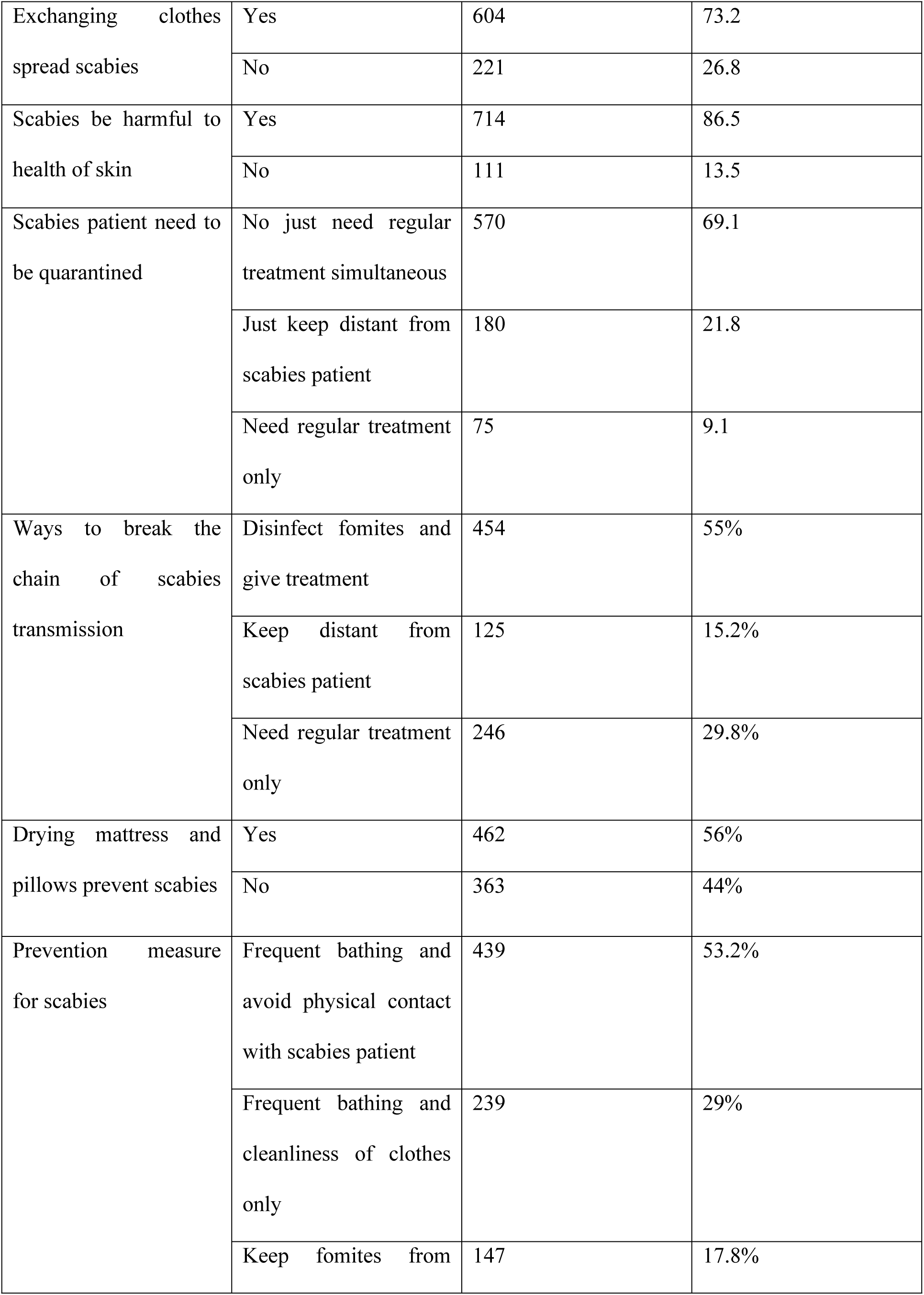
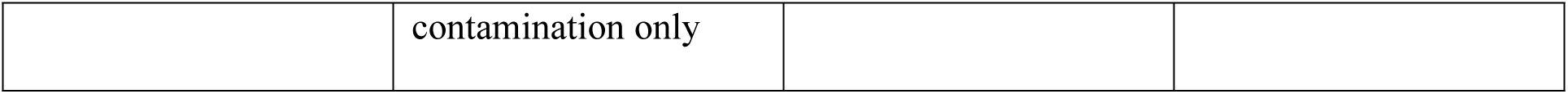
knowledge of respondents about scabies infection in the study area, 2018.

### Factors associated with scabies disease

In multivariable logistic regression analysis, overcrowding index, family member/entourage complaining of itching in the past two weeks, wealth index, knowledge of about scabies, climatic zone, frequency of washing body, frequency of washing clothes, not cutting finger nails short/trimmed, history of skin contact with scabies patient, washing hairs more than seven days and sharing of clothes were significantly associated scabies disease. Overcrowding index more than 1.5[AOR = 5.433, 95%CI: 2.264, 13.04] were 5 times more likely to have scabies than whose overcrowding index less than or equals to 1.5. Those children whose family member or entourage complaining of itching in the past two weeks [AOR = 6.99, 95%CI: 2.81, 17.406] were 7 time high risk of getting scabies when compared to no complaining in the family about itching in the past two weeks. Families who had fourth wealth quintile (AOR = 5.542, 95%CI: 1.402, 12.91) were 5.5 times more risk when compared to wealth quintile highest. Children whose family had poor knowledge about scabies [AOR = 5.2, 95% CI: 2.188, 12.358] were 5 more likely to be affected by scabies disease than those whose family had good knowledge. This study also revealed that children who were living in the lowland [AOR= 0.306, 95%CI = 0.109, 0.588] 69.4% reduced risk for scabies when compared to highland whereas children who were living in midlands [AOR = 0.053, 95%CI: 0.012, 0.24] were 94.7% less prone to scabies as compared to children living in highlands.

Those children who wash their clothes infrequently were 3.5 times higher risk of having scabies when compared to those children washing their clothes frequently [AOR = 3.53, 95%CI: 1.454, 8.566]. Children who wash their body infrequently had 6 times more prone to scabies than children who frequently wash their body [AOR = 6.321, 95%CI: 2.312, 17.284]. This study identified that children who share their clothes with others were 6 timely more likely to develop scabies than children who did not share clothes [AOR = 6.013, 95%CI: 2.51, 14.4].

This study identified that children who had history of contact were 10 times more prone to scabies than who did not had contact history(AOR = 9.579, 95% CI: 4.03, 17.22). Those children who wash their hair 7-14 days and >14 days (AOR=7.118, 95%CI: 2.63, 19.268) (AOR= 5.11, 95%CI: 1.38, 18.899) were 7 and 5 times more likely to be affected than those children who wash hair 1-7 days respectively. Children who did not cut their finger short/trimmed were 7.6 times more prone to scabies when compared to children who cut short/trimmed (AOR = 7.6, 95%CI: 3.169, 18.245) (table 6).

**Table 6:** Multivariable logistic regression analysis of factors associated with scabies among school age children in Arba Minch zuria district, 2018.

| Variables | Categories | Scabies |  |  | COR [95% CI] | AOR [95%CI] |
| --- | --- | --- | --- | --- | --- | --- |
|  |  | Yes | No | Total |  |  |
| Overcrowding Index | <=1.5 | 42[8.1%] | 476[91.9%] | 518 | 1 |  |
|  | >1.5 | 93[30.3%] | 214[69.7%] | 307 | 4.925[3.306, 7.336] | 5.433[2.264,13.04]**<br>* |
| Finger nails cut short/trimmed | Yes | 24[4.3%] | 539[95.7%] | 563 | 1 |  |
|  | No | 111[42.4 %] | 151[57.6%] | 262 | 16.509[10.249, 26.592] | 7.6[3.169, 18.245]* |
| Family member/entourage complaining of itching in the past two weeks | Yes | 83[45.4%] | 100[54.6%] | 183 | 9.417[6.273, 14.138] | 6.99[2.81,17.406]*** |
|  | No | 52[8.1%] | 590[91.9%] | 642 | 1 |  |
| Knowledge status | Good knowledge | 48[17.5%] | 588[92.5%] | 636 | 1 |  |
|  | Poor knowledge | 87[46%] | 102[54%] | 189 | 10.449[6.932, 15.749] | 5.20[2.188,12.358]**<br>* |
| Have you infected by scabies before? | Yes | 24[50%] | 24[50%] | 48 | 6[3.292, 10.937] | 0.921[0.17, 2.314] |
|  | No | 111[14.3%] | 666[85.7%] | 777 | 1 |  |
| Climatic zone | Lowland | 45[13.8%] | 280[86.2%] | 325 | 0.379[0.216, 0.667] | 0.306[0.109, 0.588]** |
|  | Midland | 65[15.6%] | 351[84.4%] | 416 | 0.437[0.255, 0.748] | 0.053[0.012, 0.24]*** |
|  | Highland | 25[29.8%] | 59[70.2%] | 84 | 1 | 1 |
| Sharing of bed with others | Yes | 122[18.5%] | 540[81.6%] | 662 | 2.607[1.431, 4.749] | 0.954[0.251, 3.635] |
|  | No | 13[7.9%] | 150[92%] | 163 | 1 | 1 |
| How often did the child change clean clothes | Frequently | 77[11.5%] | 590[88.5%] | 712 | 1 | 1 |
|  | Infrequently | 58[36.7%] | 100[63.3%] | 113 | 4.444[2.975, 6.639] | 0.567[0.174, 1.849] |
| How often did you wash your clothes | Frequently | 43[6.5%] | 619[93.5] | 662 | 1 |  |
|  | Infrequently | 92[56.4%] | 71[43.6%] | 163 | 18.653[12.043, 28.890] | 3.53[1.454, 8.566]* |
| How often did you wash your body | Frequently | 61[8.6%] | 647[91.4%] | 708 | 1 |  |
|  | Infrequently | 74[63.2%] | 43[36.8%] | 117 | 18.253[11.541, 28.869] | 6.321[2.312, 17.284]*<br>* |
| Sharing of clothes | No | 33[5.1%] | 608[94.9%] | 641 | 1 | 1 |
|  | Yes | 102[55.4%] | 82[44.6%] | 184 | 22.918[14.537, 36.131] | 6.013[2.51, 14.4]*** |
| Distance to fetch water | Near home | 100[13.4%] | 647[86.6%] | 747 | 1 | 1 |
|  | Far away | 35[44.9%] | 43[55.1%] | 78 | 5.266[3.215, 8.626] | 3.106[0.674, 14.313] |
| History of contact with scabies patient | No | 18[3.1%] | 561[96.9%] | 579 | 1 | 1 |
|  | Yes | 117[47.6%] | 129[52.4%] | 246 | 28.267[13.307, 48.114] | 9.579[4.03, 17.22]*** |
| How often did you wash hair | 1-7 day | 17[3.6%] | 451[65.4%] | 468 | 1 | 1 |
|  | 7-14 day | 81[28.9%] | 199[71.1%] | 280 | 10.798[6.237, 18.694] | 7.118[2.63, 19.268]** |
|  | >14 day | 37[48.1%] | 40[51.9%] | 77 | 24.54[12.695, 47.434] | 5.11[1.38, 18.899]** |
| Wealth Index | Lowest | 27[17.3%] | 129[82.7%] | 156 | 1.408[0.765, 2.592] | 0.571[0.146, 2.237] |
|  | Second | 23[12.9%] | 155[87.1%] | 178 | 0.998[0.534, 1.868] | 2.01[0.457, 8.87] |
|  | Middle | 25[15.8%] | 133[84.2%] | 158 | 1.265[0.681, 2.348] | 1.48[0.36, 6.072] |
|  | Fourth | 38[23.3%] | 125[76.7%] | 163 | 2.045[1.149, 3.64] | 5.542[1.402, 12.9]* |
|  | Highest | 22[12.9%] | 148[87.1%] | 170 | 1 | 1 |

P<0.05\* p<0.01\*\* p<0.001\*

## Discussion

This study was held to assess the prevalence of scabies and to determine possible risk factors among school age children. Scabies affects children life such as reducing school attendance due to severe itching. Secondary bacterial complications were another problem that affects scabies patients. This study showed that the overall prevalence of scabies was 16.4%. It is comparable with study conducted in India wardha district 18% and with a report from Cameroon 17.8% [11]. However it is much lower than study done at West Bengal India, and Pakistan with prevalence of 42% and 47.6% respectively (12, 13). This study also revealed that the prevalence is lower from the study done in Vanuatu, Solomon Island, Sierra Leon displacement camp and Malaysian welfare home with prevalence of 24%, 25%, 86% and 31% respectively [9, 12-14]. This difference might be due to difference in socio-economic characteristics between the study areas. Children who live in lowland area were 69.4% reduced risk for scabies when compared to highland whereas children who live in midlands were 94.7% less prone to scabies as compared to children living in highlands. Climatic condition was significantly associated with scabies infection. This might be explained by weather change or reduced atmospheric temperature in highland area lead to reduce the frequency of bathing of their clothes as well as their body. This finding was consistent with study done in Iran and by Hosseini-Shokouh et al [15, 16].

Evidence suggests that overcrowding was one of the main risk factor for scabies infection in different parts of the world including Ethiopia [17-21]. This study also revealed that overcrowding index was positively associated with scabies; overcrowding index more than 1.5 was 5 times more likely to scabies when compared to overcrowding index 1.5 and less. This might be due to frequent body contact with scabies patients and sharing of clothes in the family.

Study conducted in Cameroon boarding school reported that scabies had no statistically significant association with finger nails cut short [22]. In contrast, this study revealed that children who did not cut their finger nails short/trimmed were 7.6 times more prone to scabies than who cut their finger short and it is statistically significant. This might be explained by finger nails can hold scabies parasites and transmit scabies disease.

This study showed that children whose family member complaining of itching in the past two weeks were 7 times significantly higher than family member not complaining of itching. This agrees with study done in Egypt [10], and Tigray regional and SNNPR Ethiopia [23, 24]. This might be due to frequent and prolonged body contact between family members sufficient to pass the mites in to others skin.

The present study revealed that children’s family who had poor knowledge about scabies were 5 times higher risk to scabies than those family who had good knowledge. This was in line with study done in Cempaka District Banjarbaru South Kalimantan [25], and in Ethiopia [17]. This might be due to the fact that families who had knowledge about the disease take care for the children and themselves from the disease as well as be treated immediately.

Regarding personal hygiene, findings from this study revealed that washing hair more than seven days, infrequent washing of clothes and infrequent washing of body were positively associated with scabies. It was true for study conducted in Pakistan, Brazil, Egypt, Amhara and Tigray region in Ethiopia; they reported that there was significant association between the factors and scabies [17, 23, 26-29]. The reason might be the respondents had less awareness about importance of personal hygiene and poor personal hygiene might be a risk factor for spread of scabies mites.

This study also revealed that sharing clothes with other person were 6 times more likely to be affected by scabies when compared to those children who did not share their clothes with others. This finding was consistent with the study done in Doga-Tembi district Tigray [23], Gojjam Amhara region [17], and Egypt [10]. This might due to scabies mites can stay out of human skin for up to 48 hours, physical transmission of the female mites through fomites like clothes possible. Children who had history of contact with scabies patient in the past two months were 10 times more likely higher when compared to no history of contact. This might be explained by scabies was one of communicable disease which can be transmitted through physical body contact from the infected person to other healthy person. Regarding wealth index, those children family who were fourth wealth quintile about 5.5 times more risk to scabies when compared to those family with highest wealth quintiles. This might be due to those family who were highest wealth quintile had good personal hygiene practices like wearing clean clothes, not sharing clothes.

This study further revealed that all children diagnosed scabies had itchy skin rash worsen at night. Body parts affected by scabies were finger webs and ulnar area, auxiliary area, genitalia, on the abdomen, on the shoulder blades, on the elbow, on the buttocks and on the lower legs. In this study seasonal variation of the diseases and confirmatory diagnosis was not addressed.

## Conclusion

In this study, the prevalence of scabies was high among school age children. Most of the children diagnosed as scabies were living in highland area followed by midland and then lowland. Overcrowding index, knowledge status of families, family member complaining of itching in the past two weeks, washing hair more than week, wealth index, infrequent washing of clothes, infrequent washing of body, history of contact with scabies patient, washing hair more than seven days and sharing of clothes were factors associated with scabies.

## Recommendation

Based on the findings of this study, the following recommendations are proposed:

✓For Zonal and Regional health bureau
  - Should strengthen scabies prevention strategies through health promotion and education about personal hygiene in the community.
✓For Woreda health and administration offices
  - In partnership with other stakeholder should mobilize the community to have good knowledge about scabies; its mode of transmission, control measures and the importance of personal hygiene in the prevention of communicable disease like scabies.
✓For Arba Minch University and Neglected tropical disease coordination office
  - Should give due attention to tackle scabies disease among school age children.

## Abbreviations

BSc: Bachelor of sciences
CI: Confidence Interval
DALY: Disability Adjusted Life Years
EFY: Ethiopian Fiscal Year
ETB: Ethiopian Birr
GAS: Group a Streptococcus
GBD: Global Burden of Diseases
GN: Glomerulonephritis
HO: Health Officer
IRB: Institute Research Board
NGOs: Non-Governmental Organizations
NTDS: Neglected Tropical Diseases
MSc: Master of Science
OPD: Outpatient Department
OR: Odds Ratio
PCA: Principal Component Analysis
PI: Principal Investigator
PSGN: Post streptococcal Glomerulonephritis
RF: Renal Failure
RHD: Rheumatic Heart Disease
SD: Standard Deviation
SNNPR: South Nations Nationalities Peoples Region
SPSS: Statistical Package for Social Sciences
SRS: Simple Random Sampling
UI: Uncertainty Interval
WHO: World Health Organization

## Declarations

### Ethical approval and consent to participate

Ethical clearance was obtained from Arba Minch University, Institute Review Committee (IRC of AMU, Ref.No. 10994/111). Permission letters were obtained from the woreda administration office and the selected kebele leaders. The children’s families were informed about the objective of the study and written consent from each respondent’s family [assent for children] from each respondent’s family was obtained. The respondent’s confidentiality was maintained. Those children and families who suffered from scabies and who developed secondary complication were referred to health facility for anti scabies medication.

## Consent for publication

Not applicable

## Availability of data and materials

The dataset analyzed during the study was available from the corresponding authors on reasonable request.

## Authors’ Contribution

**CC**: Involved in generating the concept of this research paper, proposal writing, designing, analysis, write-up, preparation of scientific paper, and read and approved manuscript **DH**: reviewed the study plan, contributed in data analysis, and read and approved manuscript. **AA**: reviewed the study plan, contributed in data analysis, and read and approved manuscript. **AT & EG**: Involved in generating the concept of this research paper, proposal writing, designing, analysis, write-up, preparation of scientific paper, and manuscript preparation.

## Acknowledgements

First and for most we would like to thanks Almighty God, the Lord of wisdom, knowledge and understanding. Our heart full gratitude also goes to the Arba Minch zuria woreda administration office and health office for their support in giving necessary information for this research work. We are grateful to Arba Minch University for administrative, technical support to conduct the study. I would like to extend my thanks to Dr Emnet Ayana for their cooperation on training data collectors on diagnosis of scabies. We would like to express our heart full thanks to data collectors, supervisors, and study participants who helped us in the data collection.

